# Hepatic FGF21 mediates tissue tolerance during bacterial inflammation by preserving cardiac function

**DOI:** 10.1101/2020.10.05.310508

**Authors:** Sarah C. Huen, Andrew Wang, Kyle Feola, Reina Desrouleaux, Harding H. Luan, Richard Hogg, Cuiling Zhang, Qing-Jun Zhang, Zhi-Ping Liu, Ruslan Medzhitov

**Author notes:** Correspondence: Sarah Huen, M.D., Ph.D., 5323 Harry Hines Blvd., Dallas, TX 75390-9041.

## Abstract

Sickness behaviors, including anorexia, are evolutionarily conserved responses to acute infections. Inflammation-induced anorexia causes dramatic metabolic changes, of which components critical to survival are unique depending on the type of inflammation. Glucose supplementation during the anorectic period induced by bacterial inflammation suppresses adaptive fasting metabolic pathways, including fibroblast growth factor-21 (FGF21), and decreases survival. Consistent with this observation, FGF21 deficient mice are more susceptible to mortality from endotoxemia and poly-bacterial peritonitis, but not viral infection. Here we report that increased circulating FGF21 during bacterial inflammation is hepatic-derived, promotes cardiac function, and is required for survival. FGF21 signaling downstream of its obligate co-receptor beta-Klotho (KLB) is required. However, mice with central nervous system or adipose-specific deletion of *Klb* do not demonstrate any difference in response to bacterial inflammation, suggesting that multiple tissues and/or a novel FGF21 target tissue are required for the full protective effect of FGF21. These data suggest that hepatic FGF21 is a novel cardioprotective factor in bacterial sepsis.

**eTOC Summary:** In response to bacterial inflammation, hepatic fibroblast growth factor 21 (FGF21), an endocrine hormone that mediates adaptive responses to metabolic stresses such as starvation, promotes survival by supporting heart function.

## Introduction

Physio-behavioral responses to infection and inflammation such as anorexia are part of a collection of sickness behaviors that are evolutionarily conserved (Murray and Murray, 1979; Adamo, 2005; Ayres and Schneider, 2009). Anorexia of acute illness has traditionally been considered a maladaptive response in the face of a presumed hyper-catabolic state. However, anorexia of infection can be adaptive depending on the organism and pathogen (Ayres and Schneider, 2009; Wang et al., 2016; Weis et al., 2017; Wang et al., 2018). We found that the effect of glucose supplementation on survival varies depending on the type of infection and was independent of pathogen load. For example, in bacterial infection, we found that glucose supplementation is detrimental and promotes mortality without impacting pathogen burden or inflammatory magnitude. Concordantly, this detrimental effect of glucose supplementation on bacterial infection is recapitulated without active pathogens by simply using lipopolysaccharide (LPS). The capacity for an organism to survive an infection requires both disease resistance, which coordinates pathogen clearance, and disease tolerance, which reflects the organism’s ability to limit tissue damage and organ dysfunction caused by the infection (Ayres and Schneider, 2012; Medzhitov et al., 2012; Soares et al., 2014). Sepsis is a life-threatening inflammatory syndrome due to dysregulated host response to an infection (Singer et al., 2016). Identifying metabolic pathways that participate in mechanisms of disease tolerance could lead to potential therapeutic targets to limit organ dysfunction and improve survival in sepsis. A major metabolic consequence of anorexia is a switch from glucose metabolism to the utilization of alternative fuels as well as activation of other fasting metabolic pathways. We found that glucose supplementation during LPS sepsis suppresses several core components of fasting metabolism, including fibroblast growth factor-21 (FGF21), an endocrine fibroblast growth factor (FGF) hormone that mediates adaptive responses to metabolic stresses including starvation (Fisher and Maratos-Flier, 2016).

FGF21 is a member of the endocrine FGF family that includes FGF15/19 and FGF23. FGF21 requires the co-receptor β-Klotho (KLB) for downstream extracellular-signal-regulated kinase (ERK) signaling via FGF receptors (Ogawa et al., 2007; Adams et al., 2012; Ding et al., 2012). FGF21 plays an important role in mediating adaptive responses to metabolic stresses. Basally, FGF21 is expressed in the liver, pancreas, and adipose tissue. Starvation increases hepatic FGF21 production, promoting ketogenesis and gluconeogenesis while inhibiting somatic growth (Potthoff et al., 2012). Over-expression of FGF21 in mice extends lifespan (Zhang et al., 2012) at the cost of female infertility and growth defects (Ding et al., 2012; Owen et al., 2013), suggesting that FGF21 functions as a trade-off in starvation adaptation to growth and reproduction.

As FGF21 functions as an adaptation hormone to metabolic stressors (Fisher and Maratos-Flier, 2016), we hypothesized that FGF21 may also act as a protective adaptation hormone to other challenges that also lead to metabolic stress, such as infection and inflammation. We recently showed that circulating levels of FGF21 increase in mice injected with LPS and that mice lacking FGF21 are more susceptible to death from endotoxemia (Wang et al., 2016). Furthermore, treatment with FGF21 initiated after induction of bacterial inflammation can prolong survival in *Fgf21*^*-/-*^ mice. Interestingly, this protective effect from sepsis appears to be specific to bacterial inflammation, as *Fgf21*^*-/-*^ mice showed no difference in susceptibility to viral inflammatory states, such as influenza infection (Wang et al., 2016). Our findings suggest that increased FGF21 levels may represent an adaptive response that is critical for surviving other organismal challenges such as bacterial infections and could represent a novel therapeutic modality in the management of bacterial sepsis. However, the origin of circulating FGF21 during bacterial inflammation and how FGF21 supports survival in bacterial infection are not well understood. Prior work suggested that the source of FGF21 during bacterial inflammation was not likely the liver, and that the mechanism of FGF21 protection was due to ketone production (Feingold et al., 2012). However, we found that *Fgf21*^*-/-*^ mice do not exhibit any defect in ketogenesis during LPS sepsis nor have any significant differences in classic inflammatory markers (Wang et al., 2016). In this study, we investigated the tissue source and the effector function of circulating FGF21 during bacterial inflammation. Here we show that during bacterial inflammation, circulating FGF21 is liver-derived, is required for survival, and maintains cardiac function.

## Results and discussion

### Circulating FGF21 during bacterial inflammation is hepatic in origin and is required for survival

Given that we previously showed that treatment with recombinant FGF21 after LPS challenge can rescue FGF21 deficient mice from LPS-induced mortality (Wang et al., 2016), we examined whether constitutive over-expression of FGF21 could protect mice from LPS-associated mortality. Surprisingly, mice with transgenic over-expression of FGF21 driven by the liver-selective apoplipoprotein E promoter (*Fgf21-Tg*) (Inagaki et al., 2007) were not protected from endotoxemia and exhibited a trend toward increased mortality after LPS challenge (**Fig 1A**). The lack of benefit of constitutively high levels of FGF21 suggests the possibility that the regulated induction and timing of FGF21 activity may be critically important in promoting survival during endotoxemia. Indeed, we found that plasma FGF21 levels did not increase until later time points after LPS challenge (**Fig 1B**). In addition, the early response to LPS resulted in a significant suppression of hepatic *Fgf21* mRNA expression (**Fig 1C**). Despite this early suppression, hepatic *Fgf21* mRNA expression was induced to similar levels as fasting at later time points after LPS challenge, concurrent with the rise in plasma FGF21 (**Fig 1D**). We next explored the mechanism of hepatic *Fgf21* expression in response to endotoxemia. Regulation of FGF21 expression and effector functions appear to be context dependent. Several highly conserved metabolic and cellular stress response pathways, including Peroxisome proliferator-activated receptor alpha (PPARa) and the integrated stress response can lead to the expression of *Fgf21*. PPARa regulates the transcription of hepatic *Fgf21* in fasting and starvation conditions (Badman et al., 2007; Inagaki et al., 2007). Similar to fasting, the upregulation of hepatic *Fgf21* and increased circulating levels of FGF21 in response to LPS challenge are also PPARa-dependent (**Fig 1E-G**). In order to determine whether the change in hepatic *Fgf21* expression resulted in the observed increase in plasma FGF21 protein levels, we generated a mouse with hepatic-specific *Fgf21* deletion (*Alb-Cre;Fgf21*^*fl/fl*^, referred to as *Fgf21*^Δ*Liv*^, **Fig 1H** with tissue specificity). *Fgf21*^Δ*Liv*^ mice failed to increase plasma FGF21 levels after LPS challenge, definitively proving that plasma FGF21 during bacterial inflammation is hepatic in origin (**Fig 1I)**.

**Figure 1.**
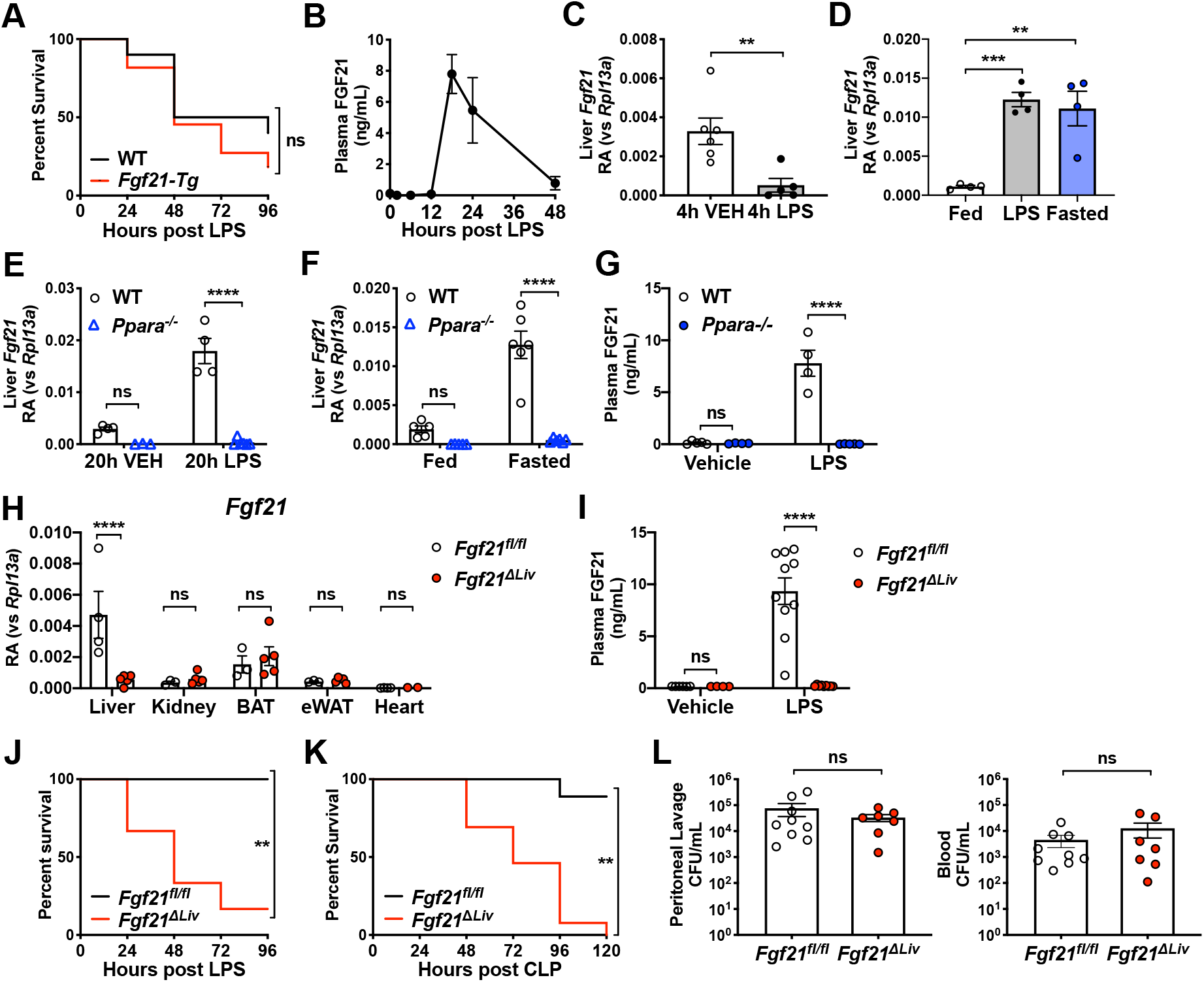
Increased circulating FGF21 during bacterial inflammation is hepatic in origin and is required for survival. (A) Kaplan-Meier survival curve after 12.5 mg/kg i.p. lipopolysaccharide (LPS) for wild-type (WT) and *Fgf21* overexpressing transgenic mice (*Fgf21-Tg*), n=10-11/group (B-C) WT mice were challenged with PBS vehicle (VEH) or 12.5 mg/kg i.p. LPS. (B) Plasma FGF21 levels measured by ELISA, n=4-5/time point. (C) Relative abundance (RA) of mRNA expression in whole liver tissue 4 hours after PBS vehicle or LPS challenge, shown relative to *Rpl13a*. n=5-6/group. (D) mRNA expression in whole liver tissue, shown relative to *Rpl13a*, 18 hours after PBS vehicle (Fed), 15 mg/kg i.p. LPS, or fasting in WT mice, n=4/group. (E) mRNA expression of whole liver tissue, shown relative to *Rpl13a*, 20 hours after PBS vehicle (VEH) or 15 mg/kg i.p. LPS in WT and *Ppara*^*-/-*^ mice. n=3-7/group. (F) mRNA expression of whole liver tissue, shown relative to *Rpl13a*, from *ad libitum* fed or 24-hour fasted WT and *Ppara*^*-/-*^ mice. n=5-6/group. (G) Plasma FGF21 levels by ELISA 18 hours after PBS vehicle or 15 mg/kg i.p. LPS in WT and *Ppara*^*-/-*^ mice. n=4-5/group. (H) Relative abundance (RA) of mRNA expression in whole tissue from *Fgf21*^*fl/fl*^ and *Alb-Cre;Fgf21*^*fl/fl*^ (*Fgf21*^Δ*Liv*^) mice, shown relative to *Rpl13a*. n=3-5/group. (I) Plasma FGF21 levels measured by ELISA 20 hours after 5 mg/kg i.p. LPS or PBS vehicle in *Fgf21*^*fl/fl*^ and *Alb-Cre;Fgf21*^*fl/fl*^ (*Fgf21*^Δ*Liv*^) mice. Vehicle n=4-6/group; LPS n=8-12/group. (J) Kaplan-Meier survival curve after 10 mg/kg i.p. LPS for *Fgf21*^*fl/fl*^ and *Fgf21*^Δ*Liv*^ mice. n=5-6/group. (K) Kaplan-Meier survival curve after cecal ligation and puncture (CLP) for *Fgf21*^*fl/fl*^ and *Fgf21*^Δ*Liv*^ mice. n=9-13/group. (L) Colony-forming units (CFUs) cultured from mouse peritoneal lavage fluid and blood 24 hours after CLP, n=7-9/group. (A,J,K) \*\**P*<0.01, ns=not significant by Log-rank (Mantel-Cox) test. (B-I,L) Data expressed as mean ± SEM. (C,L) \*\**P*<0.01, ns=not significant by two-sided, unpaired t-test. (D) \*\**P*<0.01, \*\*\**P*<0.001 by one-way ANOVA with Dunnett’s multiple comparisons test. (E-I) **\*\*\*\****P*<0.0001, ns=not significant by two-way ANOVA with Sidak’s multiple comparisons test.

In order to determine the significance of hepatic FGF21 production during bacterial inflammation, we next investigated the effect of hepatic deletion of *Fgf21* on the survival of mice during endotoxemia. Similar to *Fgf21*^*-/-*^ mice (Wang et al., 2016), *Fgf21*^Δ*Liv*^ mice were also more susceptible to mortality from LPS (**Fig 1J**), suggesting that hepatic FGF21 production induced by endotoxemia is required for survival during acute bacterial inflammation. As LPS challenge models the sterile inflammation that accompanies bacterial infection, the increased mortality in FGF21 deficient mice in the absence of live bacteria suggests that FGF21 supports survival through promoting disease tolerance. In order to determine whether FGF21 contributes to disease resistance, we employed cecal ligation and puncture (CLP), a poly-bacterial infection model. While *Fgf21*^Δ*Liv*^ mice were more susceptible to mortality after CLP (**Fig 1K**), the bacterial load was no different between *Fgf21*^Δ*Liv*^ and wild-type controls (**Fig 1L**), suggesting that hepatic FGF21 confers protection against bacterial inflammation through promoting disease tolerance, rather than via bacterial clearance.

We next examined the effect of FGF21 deficiency on physiologic and metabolic parameters in response to bacterial inflammation. FGF21 is known to promote thermogenesis in cold adaptation (Fisher et al., 2012; Ameka et al., 2019) and increase energy expenditure (Owen et al., 2014). Consistent with these known functions of FGF21, we found that FGF21-deficient mice also exhibited decreased body temperatures after CLP and after both high and low dose LPS challenge (**Fig 2A**). FGF21 also plays a major role in glucose and lipid metabolism, interacting with several metabolic mediators, including adiponectin, glucocorticoids, and thyroid hormone (Potthoff et al., 2009; Holland et al., 2013; Domouzoglou et al., 2014; Liang et al., 2014; Patel et al., 2015). During bacterial sepsis, multiple metabolic changes occur, including dysglycemia, hyperlipidemia, dysthermia and sick euthyroid syndrome (Van Wyngene et al., 2018). We reasoned that if FGF21 modulates metabolic pathways, it may play an important role in limiting metabolic derangement or supporting protective metabolic pathways in bacterial sepsis. However, we found no difference in blood glucose, free fatty acids, adiponectin, corticosterone, or thyroid hormone levels in LPS sepsis between wild-type and *Fgf21*^-/-^ mice (**Fig 2B-E, Fig S1A,B**).

**Figure 2.**
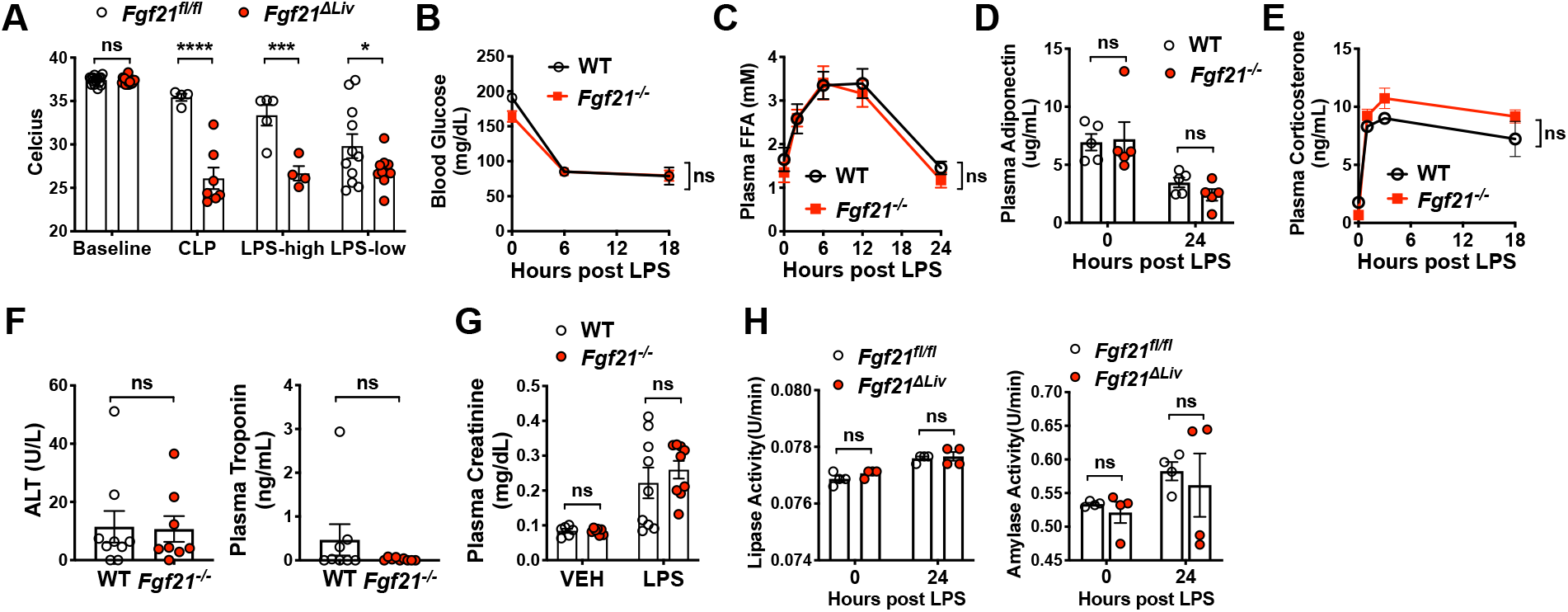
FGF21 deficiency results in decreased body temperature after LPS challenge. (A) Rectal temperatures of *Fgf21*^*fl/fl*^ and *Alb-Cre;Fgf21*^*fl/fl*^ (*Fgf21*^Δ*Liv*^) mice 24 hours after CLP, LPS-high dose (10 mg/kg i.p.) and LPS-low dose (5 mg/kg i.p.). n=4-11/group. (B-F) WT and *Fgf21*^*-/-*^ mice were challenged with 12.5 mg/kg i.p. LPS. (B) Blood glucose. n=5/group. (C) Plasma free fatty acids (FFA) by enzymatic assay. n=5/group. (D) Plasma adiponectin measured by ELISA. n=5/group. (E) Plasma corticosterone measured by ELISA. n=3-4/group. (F) Alanine transaminase (ALT) activity measured by enzymatic activity assay and troponin measured by ELISA 24 hours after LPS. n=8-9/group. (G) Plasma creatinine measured by HPLC 24 hours after PBS vehicle (VEH) or 10 mg/kg i.p. LPS in WT and *Fgf21*^*-/-*^ mice, n=6-9/group. (H) Plasma lipase and amylase activity measured by enzymatic assay before and after LPS 5 mg/kg i.p. challenge. (A-G) Data expressed as mean ± SEM. (A-E,G,H) \**P*<0.05, \*\*\**P*=0.001, \*\*\*\**P*<0.0001, ns=not significant by two-way ANOVA with Sidak’s multiple comparisons test. (F) ns=not significant by unpaired two-sided t-test. Data expressed as mean ± SEM.

We previously reported that the increased mortality in FGF21 deficient mice is not associated with increased inflammation (Wang et al., 2016). We next examined injury biomarkers of organs known to be susceptible to injury during bacterial sepsis. Interestingly, we found that FGF21 deficient animals did not exhibit significant differences in liver injury (similar alanine aminotransferase, ALT, activity), heart ischemia (normal troponin levels), or kidney injury (similar creatinine levels), compared to wild-type controls during endotoxemia (**Fig 2F,G)**. Other known biomarkers for severity of illness in sepsis including blood lactate, acidosis and hypoxia were also measured and found to not significantly differ between wild-type and FGF21 deficient animals (**Fig S1C,D**). Although the pancreas is not often an organ injured in sepsis, the clinical syndrome of pancreatitis is similar to sepsis. Furthermore, FGF21 is an important factor regulating pancreatic exocrine function and pharmacologic treatment with FGF21 can limit injury from pancreatitis (Coate et al., 2017; Hernandez et al., 2020). We found plasma amylase and lipase activity before and after LPS challenge were no different in FGF21 deficient mice compared to littermate controls (**Fig 2H**), suggesting that the protective effect of FGF21 in endotoxemia is unlikely to involve the pancreas.

### Beta-Klotho is required for survival in endotoxemia

As FGF21 signaling downstream of FGF receptors requires the co-receptor KLB (Ding et al., 2012), we next examined whether KLB is necessary for the protective effect of FGF21 in bacterial inflammation using whole body *Klb* knockout (*Klb*^*-/-*^) mice. Indeed, we found that *Klb*^*-/-*^ mice were more susceptible to mortality from endotoxemia and exhibited lower body temperatures after LPS challenge compared to heterozygous *Klb*^*+/-*^ controls (**Fig 3A,B**). As FGF21 deficient and KLB deficient animals are unable to maintain body temperature during endotoxemia, we next tested whether FGF21 was acting on the adipose tissue or centrally in the brain. In order to test the effect of FGF21 on the adipose tissue, we generated a mouse model of adipose-specific deletion of *Klb* (*Adiponectin-Cre;Klb*^*fl/fl*^, *Klb*^*ΔAdipo*^). *Klb*^*ΔAdipo*^ mice had no difference in mortality after LPS, nor had any difference in body temperature compared to *Klb*^*fl/fl*^ littermate controls (**Fig 3C,D**). While FGF21 may have a direct effect on adipose tissue for thermogenesis (Fisher et al., 2012), there are other non-adipose effects that are independent of the adipose and are likely to be mediated via the central nervous system in the context of acute cold exposure and pharmacologic dosing (Owen et al., 2014; BonDurant et al., 2017; Ameka et al., 2019). Many of the central effects of FGF21, including its effects on female reproduction, circadian behavior, metabolism, thirst response to ketogenic diet and alcohol, sweet and alcohol preference, and food choice during protein restriction, have been reported to act via KLB expression in CamK2A-expressing neurons, mainly in the suprachiasmatic nucleus (SCN) of the hypothalamus, in studies using the *Camk2a-Cre;Klb*^*fl/fl*^ mouse model (*Klb*^*ΔCamk2a*^) (Bookout et al., 2013; Owen et al., 2013; Owen et al., 2014; Talukdar et al., 2016; Song et al., 2018; Hill et al., 2019). Surprisingly, the *Klb*^*ΔCamk2a*^ mice exhibited no difference in body temperature or mortality in response to LPS challenge compared to *Klb*^*fl/fl*^ littermate control mice (**Fig 3E,F**). While liver KLB expression is high, downstream FGFR signaling via KLB in the liver is thought to be primarily activated by FGF15 (human ortholog FGF19) due to its preferential affinity for FGFR4 which is expressed in the liver (Kurosu et al., 2007). As it is unclear whether in other contexts, such as bacterial sepsis, FGF21 could signal via liver KLB, we generated animals with hepatic-specific deletion of *Klb* (*AlbCre-Klb*^*fl/fl*^; *Klb*^Δ*Liv*^).

**Figure 3.**
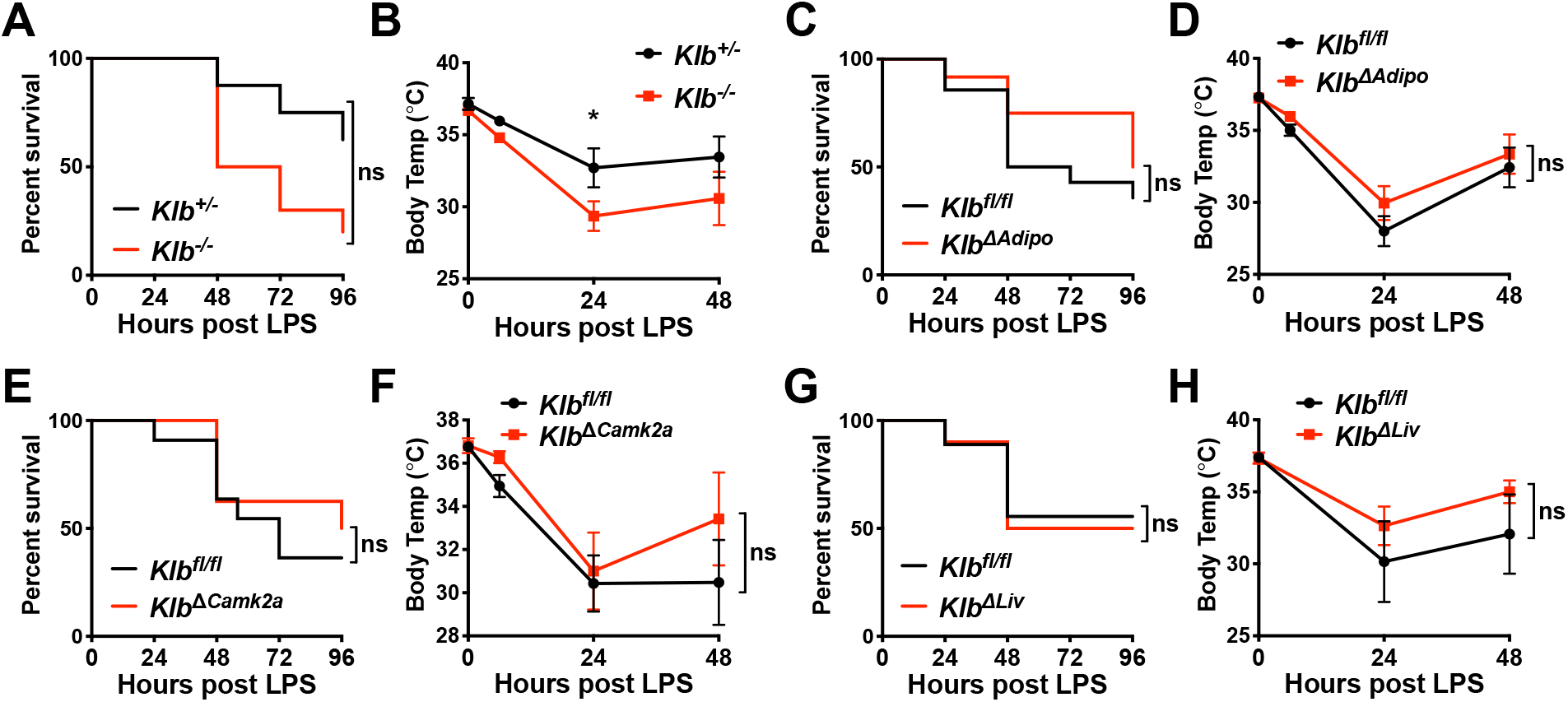
β-Klotho signaling is required for signaling for surviving endotoxemia. (A-B) *Klb*^*+/-*^ and *Klb*^*-/-*^ mice were challenged with 10 mg/kg i.p. LPS, n=8-10/group. (A) Kaplan-Meier survival curve. (B) Rectal temperatures (C-D) *Klb*^*fl/fl*^ and *Adiponectin-Cre;Klb*^*fl/fl*^ (*Klb*^*ΔAdipo*^) mice were challenged with 15 mg/kg i.p. LPS, n=10-14/group. (C) Kaplan-Meier survival curve. (D) Rectal temperatures. (E-F) *Klb*^*fl/fl*^ and *Camk2a-Cre;Klb*^*fl/fl*^ (*Klb*^*ΔCamk2a*^) mice were challenged with 15 mg/kg i.p. LPS, n=8-11/group. (E) Kaplan-Meier survival curve. (F) Rectal temperatures. (G-H) *Klb*^*fl/fl*^ and *Alb-Cre;Klb*^*fl/fl*^ (*Klb*^Δ*Liv*^) mice were challenged with 10 mg/kg i.p. LPS, n=9-10/group. (G) Kaplan-Meier survival curve. (H) Rectal temperatures. (A,C,E,G) \**P*<0.05, ns=not significant by Log-rank (Mantel-Cox) test. (B,D,F,H) \**P*<0.05, ns=not significant by two-way ANOVA with Sidak’s multiple comparisons test. Data expressed as mean ± SEM.

Similar to the brain and adipose specific *Klb* deletion mouse models, *Klb*^Δ*Liv*^ mice had similar survival and body temperature in response to LPS compared to their wild-type *Klb*^*fl/fl*^ littermates (**Fig 3G,H**). These data suggest that in bacterial sepsis, KLB is required for survival, however, the mechanism by which FGF21 acts to promote survival is either independent of the well-described FGF21 target tissues of the brain, adipose, and liver or involves FGF21 signaling on multiple KLB-expressing tissues.

### FGF21 deficiency increases cardiac dysfunction in LPS endotoxemia

Sepsis physiology is characterized by hemodynamic instability driven by reduced systemic vascular resistance and cardiac dysfunction (i.e. septic cardiomyopathy) which leads to shock and organ dysfunction (Drosatos et al., 2015). Furthermore, as a result of shunting of blood and decreased cardiac output, low body temperature could be reflective of severe heart failure (Casscells et al., 2005), often predicting poor outcomes including mortality. Moreover, recent studies of cardiomyopathy and cardiac ischemia suggest that FGF21 can act on the heart to limit cardiac injury (Planavila et al., 2013; Patel et al., 2014). We thus considered the possibility that FGF21 may affect cardiac contractility and output during bacterial sepsis by measuring cardiac function by echocardiography. *Fgf21*^Δ*Liv*^ and littermate control mice underwent echocardiograms at baseline and again 18 hours after LPS challenge, during the time when plasma FGF21 peaks (**Fig 1B**). Strikingly, despite normal troponin levels, *Fgf21*^Δ*Liv*^ mice have decreased heart rate and cardiac contractility suggesting that the LPS-induced mortality in FGF21 deficient mice may be due to cardiac dysfunction (**Fig 4A-C**). Ambulatory telemetry confirmed the decreased heart rate in *Fgf21*^Δ*Liv*^ mice after LPS observed by echocardiography, however, there was no significant difference in blood pressure in *Fgf21*^Δ*Liv*^ and littermate control mice (**Fig 4D, and Fig S2A)**. FGF21 deficiency also had no overt effect on markers of cardiac inflammation, fatty acid oxidation, and acute stress response pathways after LPS challenge (**Fig S3**).

**Figure 4.**
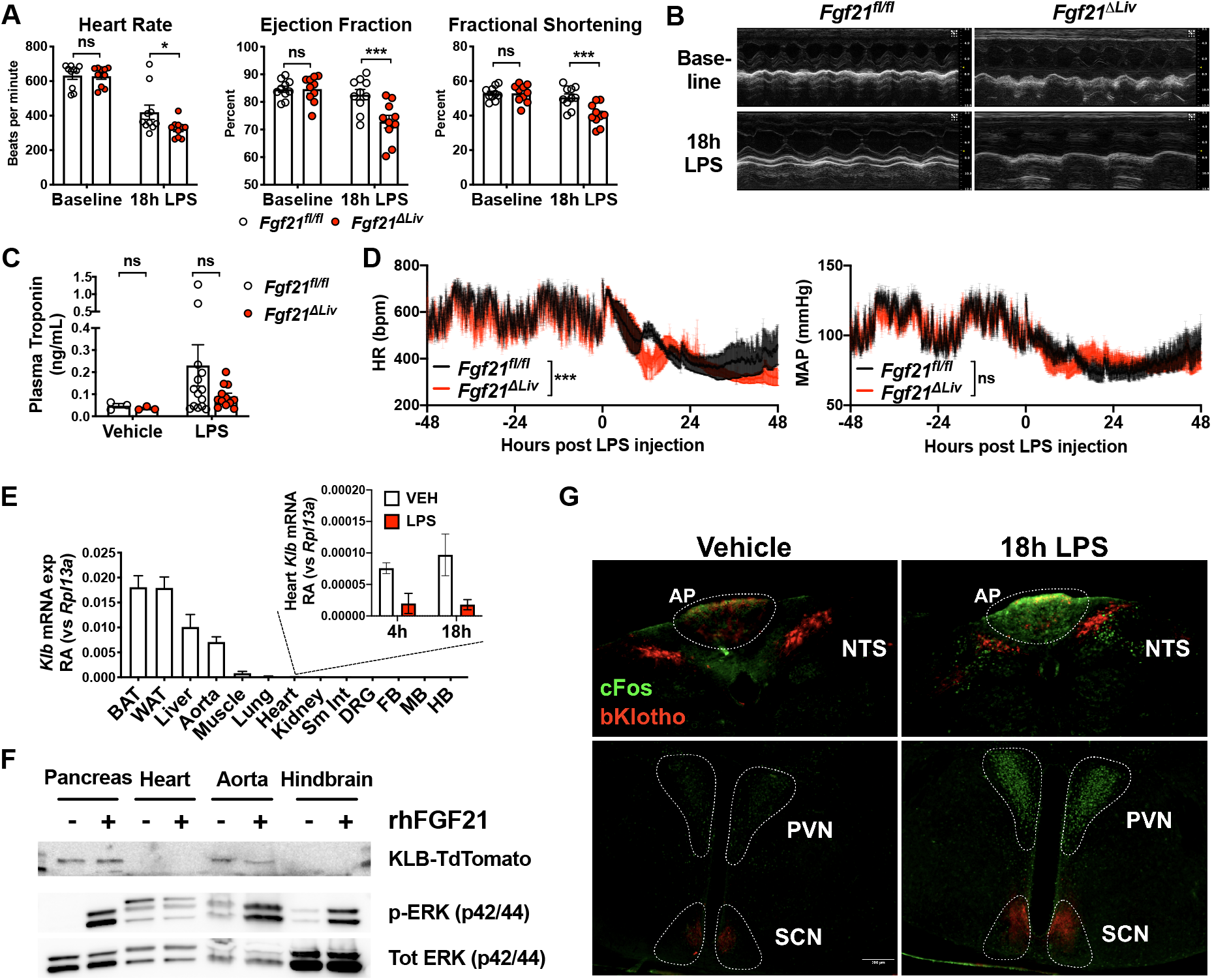
FGF21 deficiency increases cardiac dysfunction in LPS endotoxemia. (A-B) Echocardiography performed on *Fgf21*^*fl/fl*^ and *Fgf21*^Δ*Liv*^ mice before (baseline) and 18 hours after 5 mg/kg i.p. LPS. n=10/group. \**P*<0.05, \*\*\**P*<0.001, ns=not significant by two-way ANOVA with Sidak’s multiple comparisons test. Data expressed as mean ± SEM. (B) Representative echocardiogram windows. (C) Plasma troponin in *Fgf21*^*fl/fl*^ and *Fgf21*^Δ*Liv*^ mice 20 hours after PBS vehicle or 5 mg/kg i.p. LPS. Vehicle n=3/group; LPS n=10-12/group. ns=not significant by two-way ANOVA with Sidak’s multiple comparisons test. Data expressed as mean ± SEM. (D) Ambulatory blood pressure and heart rate (HR) of *Fgf21*^*fl/fl*^ and *Fgf21*^Δ*Liv*^ mice measured by *in vivo* telemetry before and after 2 mg/kg i.p. LPS. \*\*\**P*<0.001 by two-way ANOVA with Sidak’s multiple comparisons test. Mean arterial pressure (MAP) and HR measured every minute and shown as mean ± SEM within each group. (E) *Klb* mRNA expression in whole tissue from WT mice, shown relative to *Rpl13a*, inset showing relative abundance (RA) of heart *Klb* after PBS vehicle or LPS treatment. n=3-4/group. qPCR Ct values: BAT 22, WAT 22, Liver 23, Aorta 25, Heart 30 at baseline, 32 after LPS. Forebrain, FB; midbrain, MB; hindbrain, HB; brown adipose tissue, BAT; white adipose tissue, WAT; small intestine, Sm Int; dorsal root ganglia, DRG. Data expressed as mean ± SEM. (F) Whole tissue protein lysates from Klb-TdTomato reporter mice (*Klb*^*TdTm*^) immunoblotted for RFP (to detect Klb-TdTomato), p-ERK, and total ERK 10 minutes after 1 mg/kg i.p. recombinant human FGF21. Hindbrain region including area postrema and nucleus of solitary tract was grossly dissected. Representative blots from 3 independent experiments. (G) Brains from *Klb*^*TdTm*^ mice were harvested 18 hours after 15 mg/kg i.p. LPS or PBS vehicle. 50-micron fixed brain vibratomed sections were immunostained for RFP and cFos. Representative images from 3 independent experiments. Area postrema, AP; nucleus of the solitary tract, NTS; paraventricular nucleus, PVN; and suprachiasmatic nucleus, SCN. Scale bar represents 200 μm.

We next asked whether FGF21 acts directly on the heart. Prior reports suggest that KLB is expressed in the heart (Liu et al., 2013; Planavila et al., 2013; Patel et al., 2014). However, we found that relative to other tissues, *Klb* expression in the heart is extremely low and is further decreased after LPS challenge (**Fig 4E**). Furthermore, acute treatment with FGF21 did not activate ERK signaling in the heart (**Fig 4F**). Instead, we found ERK signaling in the aorta and the hindbrain (**Fig 4F**). Using a KLB-TdTomato (*Klb*^*TdTm*^) reporter mouse (Coate et al., 2017), we found KLB expression in the aorta by western blot. Similar to prior reports of *in situ* hybridization of *Klb* in the brain (Bookout et al., 2013), we found a subset of KLB-expressing cells in the SCN, area postrema (AP) and the nucleus of the solitary tract (NTS) by immunostaining (**Fig 4G**). Consistent with a lack of phenotype in the *Klb*^*ΔCamk2a*^ mice in which *Klb* is deleted from the SCN, we found that cFos, a marker of neuron activation, co-localized with KLB in the AP and NTS, but not with KLB-expressing cells in the SCN after LPS challenge (**Fig 4G**). Moreover, the *Klb*^*ΔCamk2a*^ mouse model does not effectively delete *Klb* in the hindbrain (Bookout et al., 2013).

These data collectively suggest that hepatic FGF21 maintains cardiac function during bacterial inflammation. The mechanism by which FGF21 acts during bacterial sepsis may involve direct FGF21 action on multiple tissues, or may involve an alternative FGF21 target tissue, including the endothelium, which can control systemic vascular tone and contribute to cardiac function via endothelial factors (Premer et al., 2019), and regions of the hindbrain, the AP and NTS, known to regulate autonomic function including cardiac chronotropy and inotropy (Ferguson, 1991; Jhamandas and Harris, 1992; Coote and Spyer, 2018). Furthermore, the AP and NTS receive many afferent inputs including baroreceptor, chemoreceptor and vagal afferents from the viscera and regulate many autonomic functions including the cardiovascular function such as heart rate and blood pressure (Cutsforth-Gregory and Benarroch, 2017). Activation of the cholinergic nervous system, including the AP and NTS regions, has been described in rodent models of sepsis and is associated with bradycardia (Fairchild et al., 2011). Consistent with published studies describing a significant decrease in heart rate variability immediately after acute endotoxin or pathogen exposure in mice (Fairchild et al., 2009), we also found that both *Fgf21*^*fl/fl*^ and *Fgf21*^Δ*Liv*^ mice exhibit a decrease in heart rate variability (**Fig S2B,C**). However, compared to wild-type animals, FGF21 deficient mice exhibit more bradycardia and increased heart rate variability. While this observation appears to be inconsistent with clinical data which suggests that decreased heart rate variability associates with poor prognostic indicator of sepsis (Ahmad et al., 2009), rodent models of acute bacterial infection with continuous electrocardiography suggest that the relationship of the risk of mortality and heart rate variability is more complex. While non-surviving mice had lower heart rate variability overall compared to survivors, non-surviving mice also exhibited periods of increased heart rate variability and bradycardia prior to death (Fairchild et al., 2011). The clinical significance of the pattern of heart rate depression in *Fgf21*^Δ*Liv*^ mice compared to wild-type mice before and after a brief of recovery approximately 24 hours after LPS challenge remains unclear but could reflect the development of potentially life-threatening bradyarrhythmias. The *in viv*o telemetry devices used in our studies are unable to report electrocardiography. Therefore, determination of heart rhythm and standard measurement of heart rate variability by R-R interval were not possible. While heart failure often results in relative tachycardia, decreased blood pressure and end organ damage due to decrease perfusion, *Fgf21*^Δ*Liv*^ mice were able to preserve their blood pressure with the lack of other end organ damage (lack of difference in liver and kidney injury biomarkers) suggesting the possibility of a primary autonomic defect resulting in a bradyarrhythmia resulting in decreased cardiac contractility and an increased risk of sudden cardiac death (Vaseghi and Shivkumar, 2008). Moreover, the absence of elevated troponin levels and other direct cardiac damage-related biomarkers in FGF21 deficient animals further support an indirect effect on the heart. Further research is needed to determine whether KLB-expressing regions in the AP and NTS participate in coordinating chronotropic and inotropic compensatory cardiac responses to bacterial inflammation. One compelling hypothesis based on our data is that the excessive bradycardia and high heart rate variability observed in FGF21 deficient mice may reflect an imbalance of parasympathetic and sympathetic activity resulting from abnormal signaling in the hindbrain.

In summary, we have identified hepatic FGF21 as a novel cardioprotective factor in bacterial inflammation. The regulation of hepatic FGF21 expression in endotoxemia is PPARa-dependent as it is in fasting. However, the target organ(s) of FGF21 effector function in bacterial inflammation is unclear. The lack of expression of KLB, the required co-receptor for FGF21, and the lack of FGF21 signaling in the heart suggest a novel indirect effect of FGF21 to maintain cardiac function during bacterial inflammation. A growing body of evidence (Wang et al., 2016; Rao et al., 2017; Weis et al., 2017) support the concept that in response to specific challenges, certain physiologic responses that have often been considered maladaptive, such as anorexia of acute illness or metabolic derangements of sepsis, are in fact coordinated defense mechanisms to promote survival and tissue protection. Our study further supports the concept that metabolic reprogramming through endocrine factors, such as FGF21, is a critical component of disease tolerance. Sepsis remains a deadly condition lacking therapeutic options. FGF21 may represent a promising therapeutic target, however, further research in understanding its dynamic expression and mechanism of action is needed.

## Materials and Methods

### Mice

All animal experiments were performed in accordance with institutional regulations after protocol review and approval by Institutional Animal Care and Use Committees at Yale University and University of Texas Southwestern Medical Center. *Fgf21*^*-/-*^ (Potthoff et al., 2009), *Fgf21*^Δ*Liv*^ (generated from *Fgf21*^*fl/fl*^ (Potthoff et al., 2009) and *Alb-Cre* from Jackson, Stock 003574), *Fgf21-Tg* (Inagaki et al., 2007), *Klb*^*-/-*^ (Ding et al., 2012), *Klb*^*TdTm*^ (Coate et al., 2017), *Klb*^*ΔAdipo*^ (Lan et al., 2017), and *Klb*^*ΔCamk2a*^ mice (Bookout et al., 2013) were generous gifts from Drs. David J. Mangelsdorf and Steve Kliewer. C57BL/6J (Stock 000664) and *Ppara*^*-/-*^ (*Ppara*^*tm1Gonz*^/J, Stock 008154) mice were purchased from Jackson Laboratories. *Klb*^*-/-*^ mice were maintained on a mixed (129Sv/C57BL/6) background as *Fgf15*^-/-^ and *Klb*^-/-^ mice are embryonic lethal on a pure C57BL/6 background (Kong et al., 2014). All other mouse strains were maintained on a C57BL/6 background.

For the LPS endotoxemia, mice were injected intraperitoneally with the indicated dose of LPS derived from *Escherichia coli* 055:B5 (Sigma-Aldrich) diluted in 100 µl PBS. LPS dosing varies across experiments due to mouse strain and LPS batch/lot differences. An approximate 50% lethal dose of LPS is determined empirically for each mouse strain and LPS batch. For experiments investigating TLR4-specific signaling, Ultrapure LPS (tlrl-pb5lps, Invivogen) was used to avoid potential TLR2 off-target activation by conventional LPS preparations. Core body temperature was measured by rectal probe thermometry (Physitemp TH-5 Thermalert).

Cecal ligation and puncture was perform similar to standard protocols (Toscano et al., 2011). Briefly, mice were anesthetized with a ketamine/xylazine (120 mg/kg; 16 mg/kg) mixture and given perioperative and postoperative buprenorphine (0.1 mg/kg) for analgesia. An approximate 1 cm midline laparotomy was performed, and the cecum was exposed. The cecum was ligated with 4-0 silk suture (Ethicon) 1 cm from the tip of the cecum and perforated through-and-through with a 25-gauge needle. The cecum was gently squeezed to extrude a small amount of fecal contents and returned back into the peritoneum. The peritoneal wall was closed using 4-0 chromic gut (CP Medical). The skin was closed with surgical glue and staples. Mice were given 1 ml of sterile saline subcutaneously and temporarily placed on a heating pad to aid in recovery.

### Quantification of bacterial load

Bacterial titers of peritoneal lavage fluid and blood from mice 24 hours after CLP were determined as previously described (Medina, 2010). Briefly, peritoneal lavage fluid (5 ml of sterile PBS) and blood were serially diluted, plated on Blood Agar (TSA with 5% Sheep Blood) plates (ThermoFisher), and incubated at 37°C for 18 hours.

### Plasma cytokine, metabolite, and tissue injury marker analysis

Blood glucose was determined by whole blood obtained by tail vein prick and measured using a glucometer (OneTouch). For other tests, whole blood was harvested from mice by retro-orbital bleeding and plasma was isolated using lithium heparin coated plasma separator tubes (BD). Plasma Troponin-I concentration (Life Diagnostics), Alanine Aminotransferase (ALT) activity (Cayman Chemical), β-hydroxybutyrate (Cayman Chemical), Adiponectin (Abcam), non-esterified free fatty acids (Wako Diagnostics), FGF21 (R&D and BioVendor), Corticosterone (Enzo), TSH and Free T4 (LSBio), and Lipase and Amylase activity (Sigma-Aldrich) were assayed using kits according to manufacturers’ protocols. Venous blood gas and lactate levels were measured using the i-STAT1 Handheld Analyzer (Abaxis, cartridges CG4+ and CG8+). Plasma creatinine was assayed using HPLC by The George M. O’Brien Kidney Center at Yale.

### RNA extraction and quantification

For tissue RNA extraction, tissues were harvested into RNA Bee RNA isolation reagent (Tel Test, Inc) and disrupted by bead homogenization in Lysing Matrix D tubes using a FastPrep-24 5G homogenizer (MP Biomedicals) or in Fisherbrand™ Pre-Filled Bead Mill Tubes using a Fisherbrand™ Bead Mill 24 Homogenizer (Fisher). RNA was extracted using the RNeasy or Direct-Zol Kits according to manufacturer’s protocol (Qiagen or Zymo Research, respectively). cDNA synthesis was performed using MMLV reverse transcriptase (Clontech) with oligo(dT) primers. qRT-PCR reactions were performed on either a CFX96/CFX384 Real-Time System (Bio-Rad) or QuantStudio 7 Flex (Applied Biosystems) using PerfeCTa SYBR Green SuperMix (Quanta Biosciences) or iTaqTM Universal SYBR Green Supermix (Bio-Rad), and transcript levels were normalized to *Rpl13a*. Primers used for qRT-PCR are cataloged in Table 1.

**Table 1.**
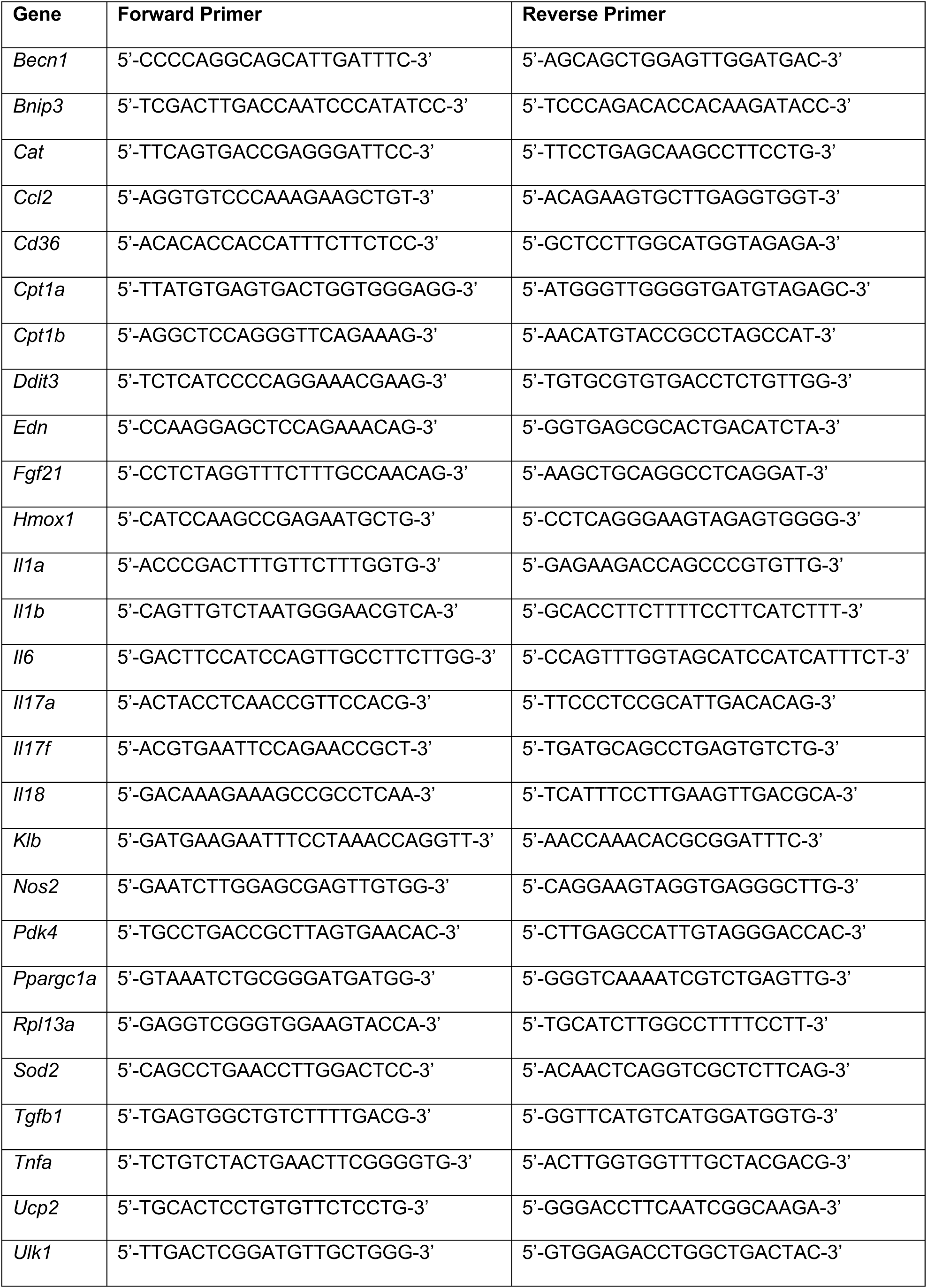

### Western Blot

Tissue samples were harvested and snap frozen in liquid nitrogen. Frozen tissues were bead homogenized in RIPA buffer (Teknova) supplemented with the HALT protease and phosphatase inhibitor cocktail (ThermoFisher Scientific), according to the manufacturer’s protocols. Equal amounts of protein were loaded per well (Bio-Rad). Protein was transferred onto activated PVDF membrane (Bio-Rad), then blocked in 5% milk in TBST (20 mM Tris, 150 mM NaCl, 0.05% Tween 20) for 30 minutes. Membranes were incubated with primary antibodies overnight at 4°C. Primary antibodies used include anti-RFP (Rockland 600401379), p-ERK (CST 4370), and total ERK (CST 4695). Membranes were washed three times with TBST, then incubated with secondary antibodies for one hour at room temperature. After washing, protein was visualized using enhanced chemiluminescence reagent (Bio-Rad).

### Trans-thoracic echocardiogram

Transthoracic echocardiography was recorded in awake mice using VisualSonics Vevo 2100 small animal echocardiography machine. Views were taken in planes that approximated the parasternal short-axis view in humans. Ejection fraction, fractional shortening, and heart rate were obtained from M-Mode parasternal short axis measurements. Ultrasound images were recorded and analyzed by an experienced technologist with expertise in mouse ultrasound imaging. The technologist was blind to the treatments and genotypes of the mice in this study.

### Ambulatory blood pressure and heart rate measurements

Experiment performed by Animal Physiology and Phenotyping Core within The George M. O’Brien Kidney Center at Yale. Briefly, a blood pressure transducer (TA11-PA-C10, DSI) was surgically implanted under anesthesia into one carotid artery. Mice were allowed to recover for 7 days prior to recording of baseline values, then given 2 mg/kg i.p. LPS. Measurements were recorded every minute.

### Immunostaining of mouse tissues

Brains from *Klb*^*TdTm*^ mice were harvested 18 hours after 15 mg/kg i.p. LPS or PBS vehicle and fixed in 10% formalin overnight. Fixed brains were then vibratomed (50 micron sections). Brain sections were immunostained for RFP (Rockland 600401379) and cFos (Santa-Cruz, (C-10) sc-271243). Images of immunostained sections were acquired using the Axio Scan.Z1 (Zeiss) and processed using Zen 2.3, Blue Edition (Zeiss).

### Statistics

Statistical information is indicated in figure legends. Statistical analyses were performed using Prism 8.0 (GraphPad Software, Inc.). Where appropriate, the Student’s t-test, ANOVA (one- or two-way) with multiple comparison analysis (Dunnett’s or Sidak’s, respectively), when appropriate. Kaplan Meier survival curves were compared using log-rank Mantel-Cox test. A *P* value less than 0.05 was considered statistically significant.

## Author contributions

S.C.H., A.W., and R.M. designed the research studies and analyzed data. S.C.H., A.W., K.F., R.D., and H.H.L. conducted experiments and acquired data with assistance from R.H. and C.Z. S.C.H. and R.M. wrote the manuscript. Q.J.Z. performed the echocardiography studies and Q.J.Z and Z.P.L. analyzed the echocardiography data.

## Acknowledgments

We thank members of the Medzhitov and the Mangelsdorf/Kliewer labs for helpful discussions and David Mangelsdorf and Steven Kliewer for making available the mouse models used in this study. This study was supported by the HHMI, Else Kröner Fresenius Foundation, The Blavatnik Family Foundation, and grants from the NIH (AI046688, AI089771, CA157461). S.C.H. was supported by NIH Grants K08DK110424, R35GM137984, and the American Society of Nephrology Carl W. Gottschalk Research Scholar Grant. A.W. was supported by NIH Grant T32 AR07107-39, K08AI128745. Plasma creatinine samples and invasive hemodynamic telemetric monitoring were performed through the George M. O’Brien Kidney Center at Yale (NIH P30-DK079310).

---

**Figure S1.**
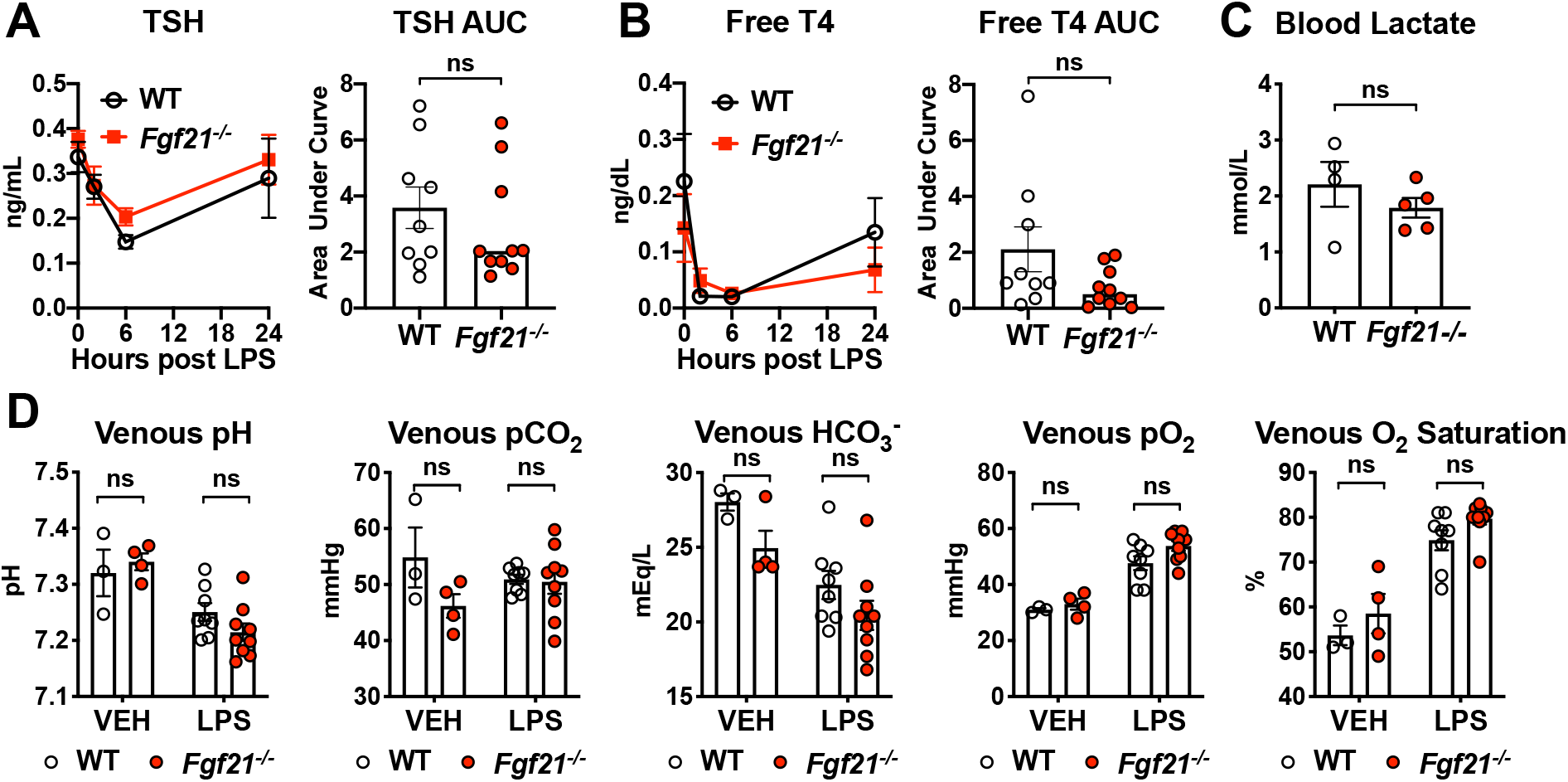
FGF21 deficient mice do not have defects in thyroid hormone axis nor differences in lactate levels or blood gases after LPS challenge. (A-B) C57BL/6J (WT) and *Fgf21*^*-/-*^ mice were challenged with LPS 10 mg/kg i.p., n=9-10/group. ns=not significant by unpaired two-sided t-test. Data expressed as mean ± SEM. (A) Thyroid stimulating hormone (TSH) measured by ELISA. (B) Free Thyroxine (T4) measured by ELISA. (C) Venous blood Lactate measured by i-STAT1 Handheld Analyzer CG4+ cartridge, 24 hours after 12.5 mg/kg i.p. LPS in WT and *Fgf21*^*-/-*^ mice, n=4-5/group. ns=not significant by unpaired two-sided t-test. Data expressed as mean ± SEM. (D) Venous blood gas measured by i-STAT 1 Handheld Analyzer CG8+ cartridge, 24 hours after PBS vehicle (VEH) or 12.5 mg/kg i.p. LPS in WT and *Fgf21*^*-/-*^ mice, n=3-9/group. ns=not significant by two-way ANOVA with Sidak’s multiple comparisons test. Data expressed as mean ± SEM.

**Figure S2.**
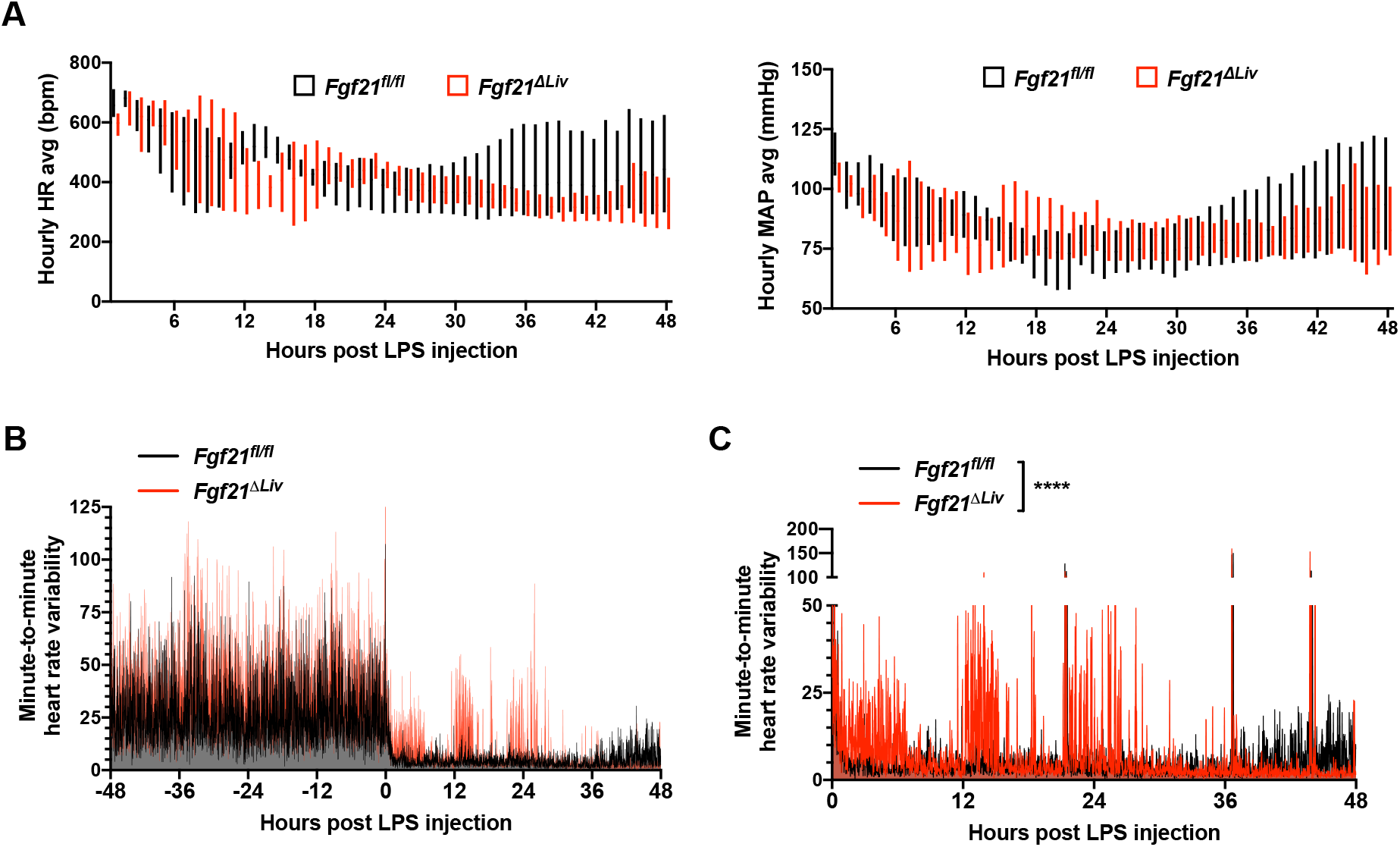
FGF21 deficiency results in depressed heart rate and increased heart rate variability during endotoxemia. Ambulatory blood pressure and heart rate measured by *in vivo* telemetry before and after 2 mg/kg i.p. LPS. Related to Main Figure 4D. (A) Hourly averages of heart rate (HR) and mean arterial pressure (MAP) after LPS injection in *Fgf21*^*fl/fl*^ vs *Fgf21*^Δ*Liv*^. Floating bars represent minimum and maximum hourly averages per genotype group. (B-C) Telemetry measurements were recorded every minute. Heart rate differences between each recording shown. (B) Minute-to-minute heart rate variability before and after LPS injection. Data expressed as mean ± SEM. (C) Minute-to-minute heart rate variability after LPS injection, *Fgf21*^Δ*Liv*^ (red curve) graphed in front to show difference. n=4/group, \*\*\*\**P*<0.0001 by two-way ANOVA with Sidak’s multiple comparisons test. Data expressed as mean ± SEM.

**Figure S3.**
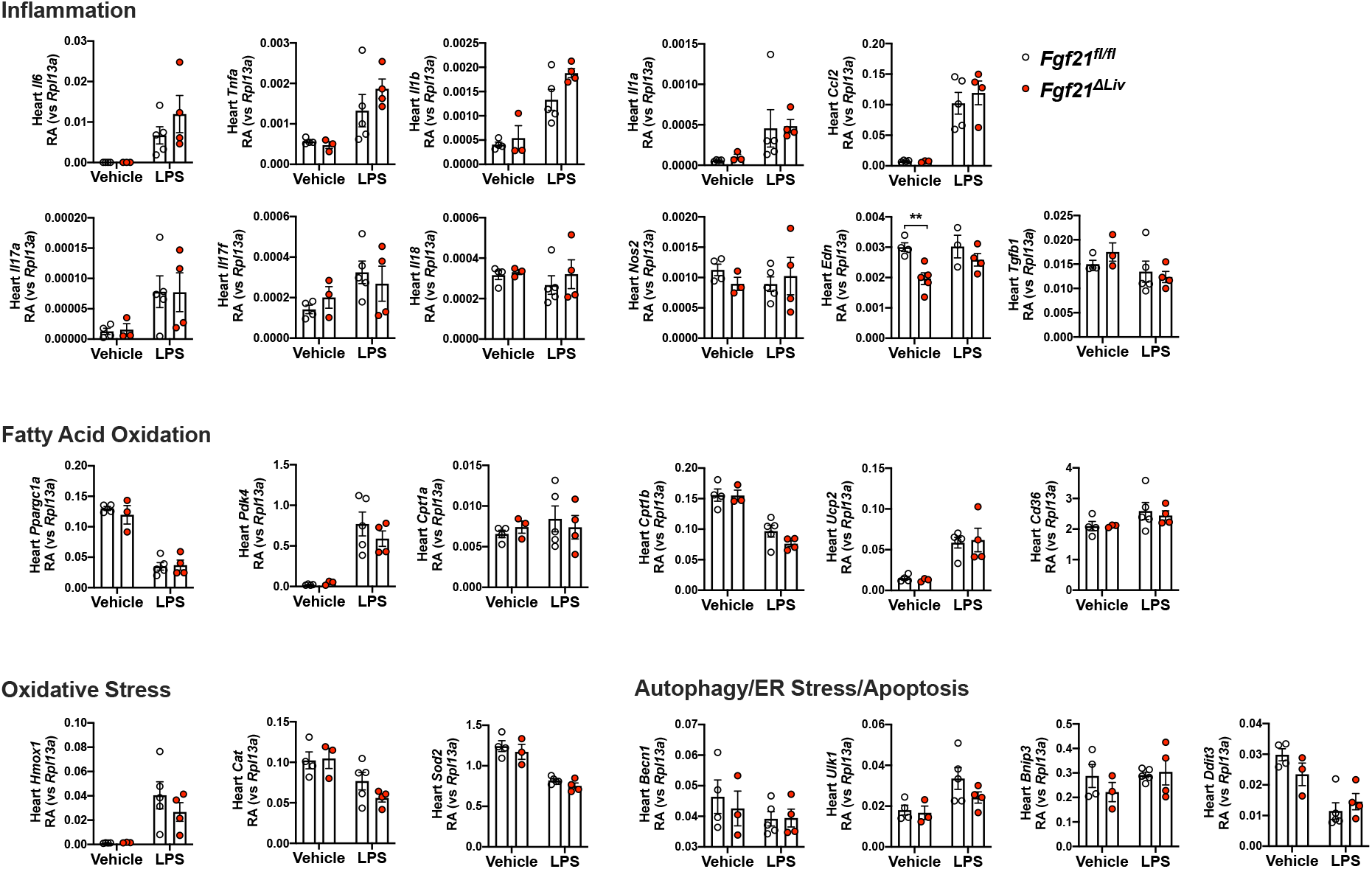
Effects of FGF21 deficiency on cardiac markers of inflammation, fatty acid oxidation, and acute stress response during endotoxemia. Relative abundance (RA) of mRNA expression in whole heart tissue from *Fgf21*^*fl/fl*^ and *Alb-Cre;Fgf21*^*fl/fl*^ (*Fgf21*^Δ*Liv*^) 20 hours after vehicle or 5 mg/kg i.p. LPS treatment, shown relative to *Rpl13a*. n=3-5/group, \*\**P*<0.01 by two-way ANOVA with Sidak’s multiple comparisons test. Data expressed as mean ± SEM

